# Compositional changes of the lung extracellular matrix in acute respiratory distress syndrome

**DOI:** 10.1101/2024.11.14.623462

**Authors:** YW Fan, J Moser, RM Jongman, T Borghuis, JM Vonk, W Timens, Meurs M van, J Pillay, JK Burgess

**Author notes:** Contributed equally (J Pillay, JK Burgess). Correspondence: Janesh Pillay, University Medical Center Groningen, Department of Critical Care, Hanzeplein 1 [IPC EA11], 9713 GZ Groningen, The Netherlands.

## Abstract

**Background:** Acute respiratory distress syndrome (ARDS) is pathologically characterized by diffuse alveolar damage (DAD) and is associated with high morbidity and mortality rates. Remodeling of the extracellular matrix (ECM), which is pivotal for both tissue repair and organ recovery, may play a large role in persistent ARDS. This study investigated the compositional changes in the ECM in different DAD stages in ARDS.

**Methods:** Paraffin-embedded lung sections collected during autopsy or from post-transplant lungs were obtained from patients with ARDS (n=28) admitted to the University Medical Center Groningen between 2010-2020. Sections were stained histochemically, and immunohistochemically for collagen III α1 chain (Col IIIa1), IV α3 chain (Col IVa3), VI α1 chain (Col VIa1), periostin (PSTN), lumican (LUM), and fibronectin (FN). The sections were divided into 118 regions based on DAD stages (54 early vs 64 advanced). The differences in the expression of selected proteins were compared between DAD stages or across ARDS duration (<7days, 7-14days, >14days). The fiber pattern of Col VIa1 was analyzed using CellProfiler.

**Results:** Higher tissue density, lower proportional areas of Col IIIa1, Col IVa3, and LUM, and more concentrated Col VIa1 fibers were observed in the advanced DAD stage than in the early DAD stage. Areas with higher proportions of total collagen and FN, and lower proportional areas of Col IIIa1, Col IVa3, and LUM were detected in lung regions from patients with ARDS >14days duration.

**Conclusions:** These findings revealed proportional changes in ECM components, strongly suggesting that dynamic changes in ECM proteins play a role in pathophysiology in ARDS during progression.

## Introduction

Acute Respiratory Distress Syndrome (ARDS) is clinically defined as the rapid onset of hypoxemic respiratory failure with non-cardiogenic bilateral pulmonary edema requiring mechanical ventilation or high-flow nasal oxygen (1). The defining pathological feature of this syndrome is diffuse alveolar damage (DAD), which describes morphologic changes specifically appearing in the lung parenchyma, involving sequentially occurring exudative, fibroproliferative, and fibrotic phases (2). The presence of DAD is considered to be the histopathological hallmark of ARDS (3–5). Patients who meet both clinical ARDS definitions and exhibit DAD on open lung biopsy have higher mortality and longer ICU stay than those who only meet the clinical ARDS definition without DAD (6, 7). These studies suggest that DAD plays an important role in ARDS progression, the persistence of respiratory failure and higher mortality risk.

DAD is characterized by severe damage to the alveolar-capillary barrier and encompasses a series of pathological changes that compromise the normal structure and function of the lung. Key features include various stage-specific pathological symptoms such as intra-alveolar edema, alveolar type I cell necrosis, proliferation of alveolar type II cells progressively covering the denuded alveolar-capillary membrane, presence of hyaline membranes, interstitial proliferation of fibroblasts and myofibroblasts, and organizing interstitial fibrosis (8, 9).

Among the chronological stages of DAD, the fibroproliferative phase marks the critical turning point between the resolution of pulmonary edema and fibrosis, and is characterized by cell proliferation and excessive extracellular matrix (ECM) deposition (10). ARDS-associated lung proliferative and fibrotic changes have been reported to be closely associated with poor outcomes, including irreversible lung fibrosis and high mortality (11–13). In a recent study, it was shown that ARDS patients with lung fibroproliferation, can be identified by measuring the ECM synthesis biomarker, the N-terminal peptide for type III procollagen (14).

The ECM is a complex network of proteins that provides structural and biochemical support to the surrounding cells and undergoes significant remodeling during ARDS (15, 16). Regulated ECM remodeling with a stable composition is important for forming a homeostatic microenvironment, as well as maintaining tissue and organ functionality (17). Compositional alterations might participate in disease development by affecting the biochemical and biomechanical properties of the lung ECM (18–21).

Although fibroproliferation and fibrotic changes associated with ARDS are widely reported (22, 23), most studies have focused on evidence of ECM remodeling in the systemic circulation or bronchoalveolar lavage fluid (BALF) (16, 24), or simply concluded the fibrotic change with a thickened alveolar wall as increased collagen deposition, without in-depth detection of the composition (2). In this study, we investigated compositional changes in the parenchymal ECM by immunohistochemistry and image analysis. By semi-quantifying the proportional changes of selected ECM proteins in human lung tissue, we provide a clearer understanding of ECM dynamics during ARDS progression in critically ill patients.

## Materials and methods

### Ethics statements

The study was conducted in accordance with the Research Code of the University Medical Centre Groningen (UMCG), as stated on https://umcgresearch.org/w/research-code-umcg as well as national ethical and professional guidelines Code of Conduct for Health Research (https://www.coreon.org/wp-content/uploads/2023/06/Code-of-Conduct-for-Health-Research-2022.pdf). The use of left-over lung tissue in this study was not subject to Medical Research Human Subjects Act in the Netherlands, as confirmed by a statement of the Medical Ethical Committee of the University Medical Centre Groningen and exempt from consent according to national laws (Dutch laws: Medical Treatment Agreement Act (WGBO) art 458 / GDPR art 9/ UAVG art 24). All donor material and clinical information were deidentified prior to experimental procedures, blinding any identifiable information to the investigators.

### Definition of ARDS and the identification of distinct histopathological phases

For inclusion in this study, patients were required to meet both the 2012 Berlin ARDS definition (25) and have archival formalin-fixed paraffin-embedded (FFPE) lung tissue blocks with identifiable diffuse alveolar damage (DAD) available for research.

In brief, the Berlin definition of ARDS includes an acute onset of hypoxic respiratory failure within one week of a known clinical insult, evidence of bilateral chest opacities, and non-cardiac respiratory failure, with a ratio of arterial oxygen partial pressure (PaO_2_) to fraction of inspired oxygen (FiO_2_) ≤ 300 mmHg with mechanical ventilation (25). Persistent ARDS was defined as ARDS that persisted over 7 days from onset.

In our study, we defined two pathological stages of DAD: early and advanced. Early DAD is characterized by extensive intra-alveolar edema in the presence of hyaline membranes. Alveolar spaces can contain fibrin-rich exudates and blood cells. Inflammation may be present depending on the etiology.

Advanced DAD was marked by the proliferation of fibroblasts and myofibroblasts in the lung alveolar interstitial septa, which in severe cases extended into the alveolar space. The lung tissue exhibited extensive interstitial collagen deposition. Such fibroproliferative changes then result in architectural changes in the alveoli, which might even resemble non-specific interstitial pneumonia. Inflammatory cell composition, if present, shifts towards mononuclear cells, such as macrophages and lymphocytes.

Within both pathological (early and advanced DAD) groups, organizing pneumonia (OP) was sometimes found to be present. OP is defined as a non-specific inflammatory condition of the lungs, characterized by the presence of fibroblast plugs within the alveolar ducts, and alveoli, occurring as a (attempted) reparative response to the widespread alveolar damage seen in ARDS.

### Lung tissue selection and processing

The lung tissue sections used for immunohistochemistry were obtained from the pathology archive at the UMCG, derived from lung tissue blocks collected following autopsy (23 donors) or explanted lungs following transplant surgery for patients with ARDS in the ICU in the UMCG (5 donors). The screening and inclusion process is shown in Figure 1. Briefly, the pathologist (WT) screened the available lung tissue blocks to identify those with DAD. Then, the intensivist (JP) checked the clinical records of those patients during the ICU stay and selected patients who met the 2012 ARDS Berlin criteria (25). From those patients who met both DAD criteria and 2012 ARDS Berlin criteria, 28 patients were randomly selected based on the length of the ICU stay (<7 days (N=10); 7-14 days (N=8); >14 days (N=10)) and a varied distribution of ARDS etiology (e.g. extrapulmonary vs pulmonary causes).

**Figure 1.**
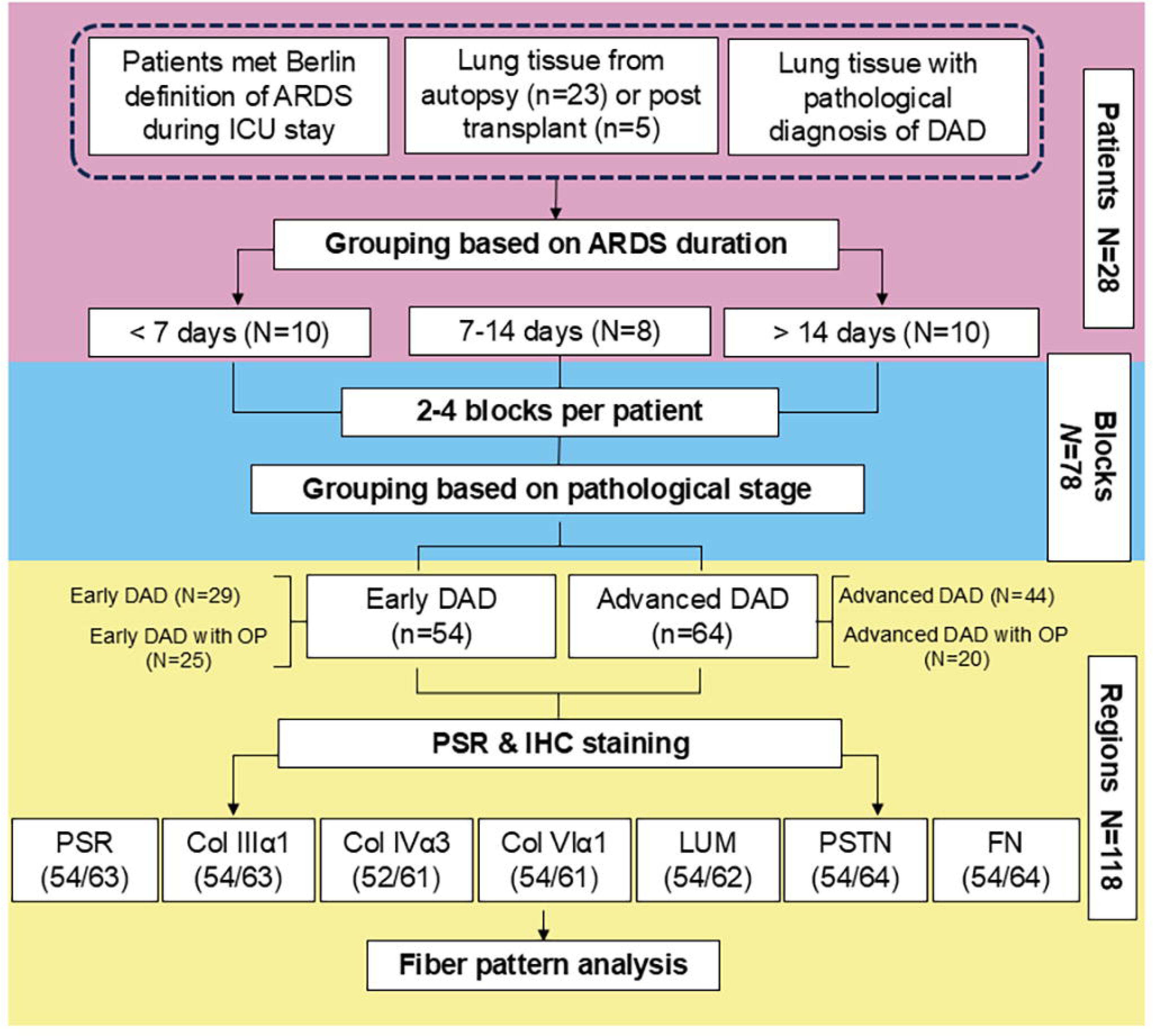
Flowchart of ARDS patients’ inclusion and blocks selection. Clinical records were screened by an intensivist, and lung tissues from the clinical archive at UMCG, from autopsy or explanted lungs following transplantation, were screened for DAD by a pathologist. A total of 28 patients were selected for this study. Patients were further categorized based on the duration of ARDS. 2-4 lung tissue blocks from each patient were used for analysis, selected blocks represented the earliest and most advanced pathological stages of ARDS. In each pathological group, regions were further divided into subgroups based on the presence or absence of OP within the tissue area. One section per block was stained for total collagen (PSR), Col3a1, Col4a3, Col6a1, LUM, PSTN, and FN before being digitally imaged and analyzed. Numbers indicate regions available for analyses (early DAD/advanced DAD). ICU, intensive care unit; DAD, diffuse alveolar damage; ARDS, acute respiratory distress syndrome; OP, organizing pneumonia; PSR, Picrosirius Red; IHC, immunohistochemistry; Col3a1, Collagen Type III α1 Chain; Col4a3, Collagen Type IVα3 Chain; Col6a1, Collagen Type VIα1 Chain; LUM, lumican; PSTN, periostin; FN, fibronectin.

For each of the 28 patients, we sought to select three blocks per patient, in which one block contained mainly parenchymal tissue, while the other two represented the earliest and advanced pathological stages of DAD, respectively. During the selection process, it was found that the same block often contained more than one of the tissue types sought, e.g. one block contained regions of tissue with both advanced DAD and also parenchymal tissue. As a result, for seven patients only two blocks were selected, which provided regions representing all the tissue types sought in our study. For one patient, four blocks were selected to ensure that all regions of interest in our study were represented. In total, 78 blocks were selected for further processing.

In the 78 blocks, regions with early and advanced pathological stages were identified and separated for subsequent analysis. Twenty blocks were found to contain areas with both early and advanced pathological stages within these blocks. Therefore, these blocks were divided into different regions to capture early and advanced DAD separately. In 10 of these 20 blocks, there were multiple separate areas within the tissue with the same pathological stage; these were captured as different regions and identified as different regions from the same block for further analysis. Any regions smaller than 1/20^th^ of the original tissue size were discarded. A total of 118 regions were collected.

The 118 regions were divided into two groups based on pathological characteristics: the early and advanced DAD groups. Each pathological group was further divided into 2 subgroups based on the presence or absence of OP within the tissue region.

#### Lung tissue chemical staining and immunohistochemistry (IHC)

FFPE blocks of lung tissue obtained from the pathology archive at the UMCG were cut into serial 4μm sections and deparaffinized, and rehydrated.

To visualize the basic anatomical structures and collagen distribution within the lung tissue, hematoxylin and eosin staining (H&E) and Masson’s trichrome staining (MTS) were performed on the lung sections, as previously described (26, 27). To visualize collagen deposition, Picrosirius Red (PSR) staining was conducted using a solution containing 0.1% PSR (PSR; Sigma-Aldrich) dissolved in 1.3% aqueous solution of picric acid (28). Staining with each chemical stain was performed in a single batch for sections from all blocks.

Specific antibodies against each protein were used for immunohistochemical staining to detect the deposition and distribution of collagen type IIIα1 (Col IIIα1), collagen type IVα3 (Col IVα3), collagen type VIα1 (Col VIα1), fibronectin (FN), lumican (LUM), and periostin (PSTN) in lung tissues. The antibodies and staining conditions are listed in Supplemental Table S1. Briefly, the sections were deparaffinized in xylene followed by a rehydration series in alcohol. Heat induced epitope retrieval or enzyme digestion was used as antigen retrieval method. After cooling down, the sections were washed with phosphate-buffered saline (PBS) before endogenous peroxidase activity was blocked using hydrogen peroxidase (0.3%) in PBS for 30 min at room temperature. After 3X PBS washes, primary antibodies diluted in 1% BSA/PBS were added to the sections and incubated overnight at 4°C (except FN; see Supplemental Table S1). Sections stained for Col IIIα1 went through an additional blocking step with incubation in 1% Bovine Serum Albumin (BSA)/PBS for 60 mins followed by overnight incubation with primary antibody.

After primary antibody incubation, the sections were washed in PBS and incubated for 45 min at room temperature with horseradish peroxidase (HRP)-conjugated goat anti-rabbit secondary antibody, which was diluted in 1% BSA-PBS solution containing 2% human serum. After washing, the sections that were stained for Col IIIα1, Col IVα3, and FN, were incubated for another 45 min with tertiary antibody HRP-conjugated rabbit-anti-goat at room temperature. Positive signal in the sections was visualized using Vector® NovaRED® substrate (SK-4800, Vector Laboratories, Canada). Hematoxylin was used for counterstaining. A negative control slide (without the primary antibody) was included in each staining batch. All sections stained for the same target protein were processed at the same time to minimize batch differences.

After dehydration with alcohol, the sections were mounted with Permount^TM^ Mounting Medium (SP15-100, Thermo Fisher Scientific, Waltham, MA, USA), and scanned at 40x using a NanoZoomer XR digital slide scanner (Hamamatsu Photonics K.K., Japan). The images were viewed and further processed using Aperio ImageScope software (v12.4.6.5003, Leica Biosystems, Germany). The quality of all stainings and images were checked to ensure technical validity; images from six slides were discarded (one from Col IIIα1, Col IVα3, Col VIα1, PSR, and two from LUM), leading to variation in the number of regions analyzed for the different proteins. The workflow for image generation and processing is shown in Figure 2.

**Figure 2.**
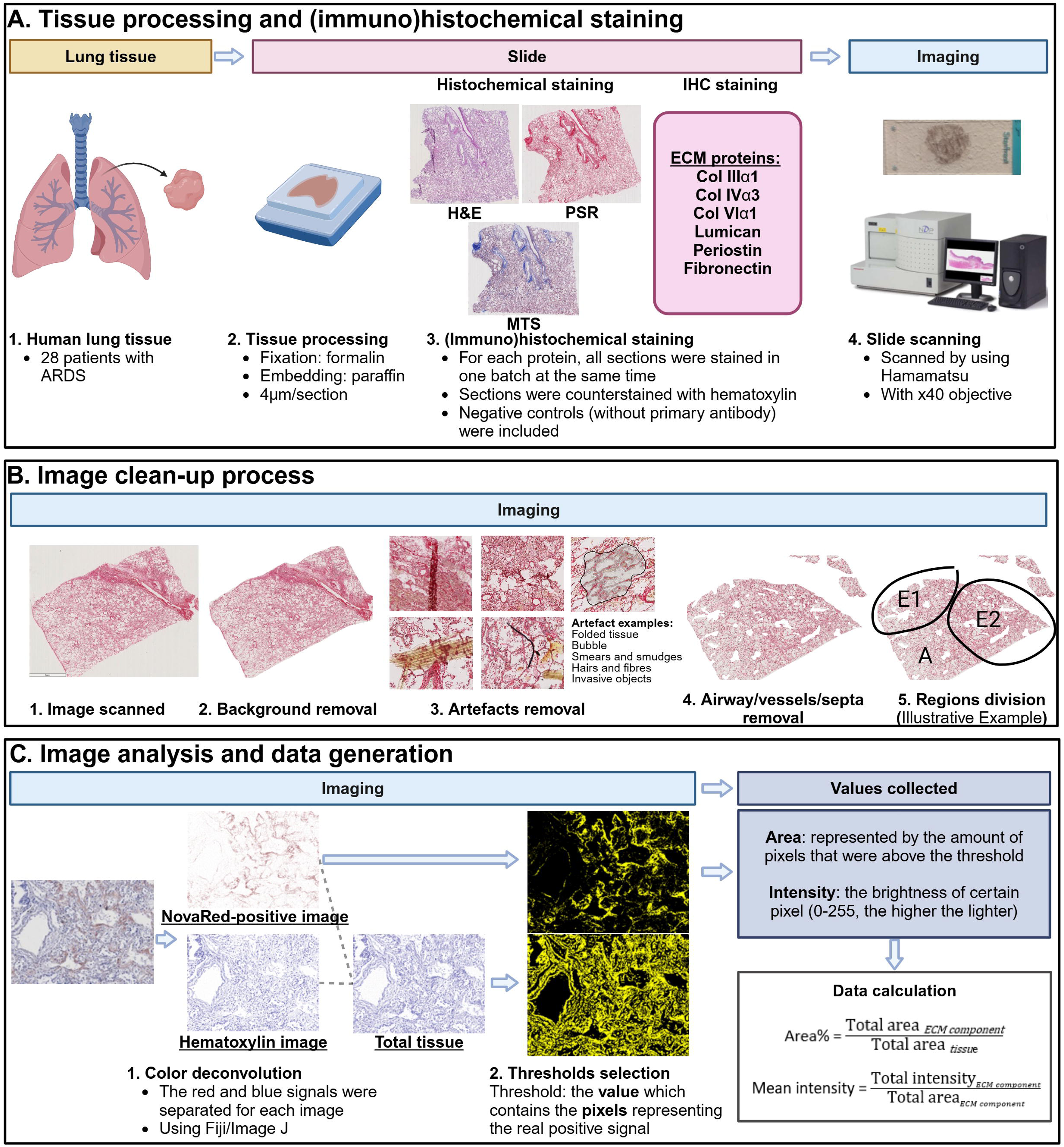
Schematic representation of the processing steps of lung tissue and digital image analyses. Representation of the workflow for processing lung tissue blocks, staining, image processing and analyses. A: Tissue processing and IHC staining; B: Image cleanup process. The clean-up process was performed using the Adobe Photoshop software to obtain the lung parenchymal area from the image. The regions in the different pathological stages (E=early phase, A=advanced phase) were divided manually; C: Image analysis. Images were separated into color layers representing the hematoxylin and NovaRed (or PSR) staining in each image. The areas from the combined image generated from the NovaRed and hematoxylin layers represented the total tissue. Color deconvolution and individual data generation were conducted using Fiji/ImageJ software. R software was used for data cleanup and further analysis. ARDS, acute respiratory distress syndrome; IHC, immunohistochemistry; H&E, hematoxylin and eosin; PSR, Picrosirius Red; MTS, Masson’s trichrome staining; Col IIIα1, collagen type 3 chain; Col VIα1, collagen type VI α Created in BioRender. Burgess, J. (2024) https://BioRender.com/l67j064.

#### Image processing and data generation

To generate data from the images, our image-processing workflow consisted of three steps (Figure 2): image extraction, clean-up, and analysis.

First, after scanning, the areas of the scans containing the lung tissue were extracted from the total scans and compressed using Aperio ImageScope software V.12.4.3 (Leica Biosystems, Nussloch, Germany).

Second, based on the anatomical characteristics of ARDS, where damage is mostly in the lung parenchymal areas, our research focused on analyzing ECM changes specifically within these areas. To obtain the specific regions of interest (parenchymal area), lung images were processed through the following steps using Adobe Photoshop software (Adobe Inc. CA) (Figure 2):

A. Background removal: Remove the empty background in the image around the tissue.
B. Artefacts removal: Remove those artefacts which do not represent the natural condition of the tissue, including the following artefacts in the parenchyma area: folded tissue; hairs and fibers; smears and smudges (regions not in focus); bubbles.
C. Airway/vessels/septa removal: Remove the visible lumen structures (airways/vessels) featuring a clear smooth muscle layer, or lung lobular and intralobular septa, both of which are rich in collagen and various extracellular matrix (ECM) components, and therefore potential confounders in our analysis strategy.

Thirdly, the processed images were manually divided into regions with the different pathological stages (E=early phase, A=advanced phase) using Adobe Photoshop software, according to the denotation by the pathologist.

Fourth, using Fiji (National Institutes of Health, Bethesda, Maryland, USA) (29), the cleaned images were deconvoluted into blue (hematoxylin-stain) and red (NovaRed or PSR stain) images using the color deconvolution plugin developed by Landini et al. (30). Optimal optical density vectors were developed individually for each ECM staining following a previously published protocol (31). The thresholds were defined to include pixels showing a positive signal and excluding the background. The images were analyzed in batches using the plugin “Slide J” in Fiji, and the details of the optimized macros used for different ECM proteins are shown in Supplemental Material S1. The numbers generated by Fiji were the total number of pixels and the total intensity of the pixels that met the criteria defined by the threshold, representing the area and positive signal level of the analyzed signal in the selected tissue, respectively. The total area of the lung tissue was calculated as the total sum of the pixels meeting the criteria for positivity in both the hematoxylin and Nova Red channels. The deconvolution vector was optimized to separate red blood cells (RBC) and other tissue staining into a background image to reduce the effect of the infiltrating RBC on the calculation of the percentage area covered by tissue.

### Col VIα1 bundle definition and detection

Cell Profiler (Broad Institute of MIT and Harvard, Cambridge, Massachusetts, USA)(32) was used to detect the specific Col VIα1 fiber pattern, referred to as “bundles” in this manuscript.

A “bundle” of Col VIα1 was defined as an aggregation of concentrated and condensed fibers, which was detected as the “object” in Cell Profiler using the following steps. The output color deconvoluted files, which contained the Nova Red signal for Col VIα1, were used as the input for the process. This process is illustrated in Supplemental figure S1.

1. The images were cut into tiles by Fiji using the plugin “Slide J,” in which the size was set to 10000 x 10000 pixels.
2. Images from regions of the same pathological stage from the same block that were smaller than 4000 × 4000 pixels were manually combined into one field of view using Photoshop. Images that did not contain any signal were discarded.
3. The processed images were first inverted (using the “invert” operation in the Cell Profiler), which reversed the intensity values of the pixels in each image, transforming the dark regions to light, and vice versa.
4. Objects in images were detected with Cell Profiler using the “Adaptive” strategy, with the “minimum Cross-Entropy” method, and the lower and upper bounds on the threshold set as “0.2-1” respectively. The typical object diameter was set to 300-2000 pixels (68.1-454µm).

The outputs captured from Cell Profiler were the “Total Area” of image, “Area Occupied” by objects and the “Count” of objects. The result was corrected for magnification to account for the unit difference between the Cell Profiler default setting (1 mm per pixel) and the input image resolution (0.227μm per pixel). The equations used to calculate the reported Col VIα1 bundle properties are as follows.

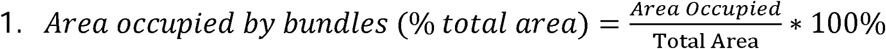

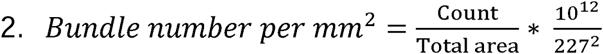

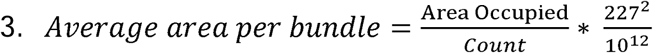

#### Data and Statistical analyses

Categorical and continuous data are presented as percentages and medians [interquartile ranges IQR], respectively.

R software was used for wrangling the data generated using Fiji. Graphs were made using GraphPad Prism version 9.1.0 (GraphPad Software, Boston, Massachusetts, USA).

Data distribution was examined using the histograms and P-P plots. Data with abnormal distributions were transformed using a natural logarithm (ln) transformation. Linear mixed model analysis with a random effect (patients and blocks) on intercept, adjusted for sex and age, was conducted to determine the ECM protein expression differences between different groups (when comparing DAD stages, the early DAD phase was used as the reference group; when comparing clinical duration of ARDS, the ARDS <7 days duration was used as the reference group). When partitioning the images by clinical duration of ARDS all regions (both early and advanced DAD) from a patient were combined to generate data for that patient. IBM SPSS Statistics for Windows (version 28.0. Armonk, NY: IBM Corp) was used for all statistical analyses.

For all tests, a *p* value of < 0.05 was considered to be statistically significant.

## Results

### Distribution regions with different DAD stages in lung tissue varies with ARDS duration

The demographic and clinical characteristics of the 28 patients included in this study are shown in Table 1.

**Table 1.** ARDS Patient characteristics.

|  | Total<br>patients | ARDS duration |  |  |
| --- | --- | --- | --- | --- |
|  |  | <7 days | 7-14 days | >14 days |
| Number, *N* | 28 | 10 | 8 | 10 |
| Age, years, median [IQR] | 56.5 [40.5-63.3] | 57.0 [43.8-64.8] | 57.0 [47.8-68.3] | 47.0 [33.0-61.0] |
| Male, *N* (%) | 16 (57.1) | 5 (50.0) | 4 (50.0) | 7 (70.0) |
| Pulmonary origin, *N* (%) | 21 (75.0) | 8 (80.0) | 6 (75.0) | 7 (70.0) |
| ARDS duration, days, median [IQR] | 11.5 [4.0-30.0] | 3.0 [2.0-4.0] | 11.5 [8.8-14.0] | 41.5 [30.0-64.8] |
| Autopsy (tissue origin), *N* (%) | 23 (82.1) | 10 (100.0) | 8 (100.0) | 5 (50.0) |
| Early DAD (regions), *N* (%) | 54 (45.8) | 28 (51.9) | 19 (35.2) | 7 (13.0) |
| Advanced DAD (regions), *N* (%) | 64 (54.2) | 12 (18.8) | 23 (35.9) | 29 (45.3) |

ARDS, acute respiratory distress syndrome; DAD, diffuse alveolar damage; IQR, interquartile range.

Lung tissues were examined by a lung pathologist and divided into regions of the early and advanced DAD stages. As expected, the greatest proportion of early DAD regions (28 of 54, 51.9%) was present in patients with an ARDS duration of less than 7 days. This proportion of early DAD regions was lower as the duration of ARDS extended. In contrast, patients with a shorter ARDS duration had fewer regions with advanced DAD. Most instances of advanced DAD were found in tissues from patients with an ARDS duration longer than 14 days (29 of 64, 45.3%). (Table 1)

### Higher proportional tissue area in the lung regions with advanced DAD pathological phase

We first examined the proportion of each lung tissue region image occupied by the tissue (mainly extracellular matrix proteins). Representative images of H&E and PSR staining from the lung regions of the early and advanced DAD groups, respectively, are presented in Figure 3.

**Figure 3.**
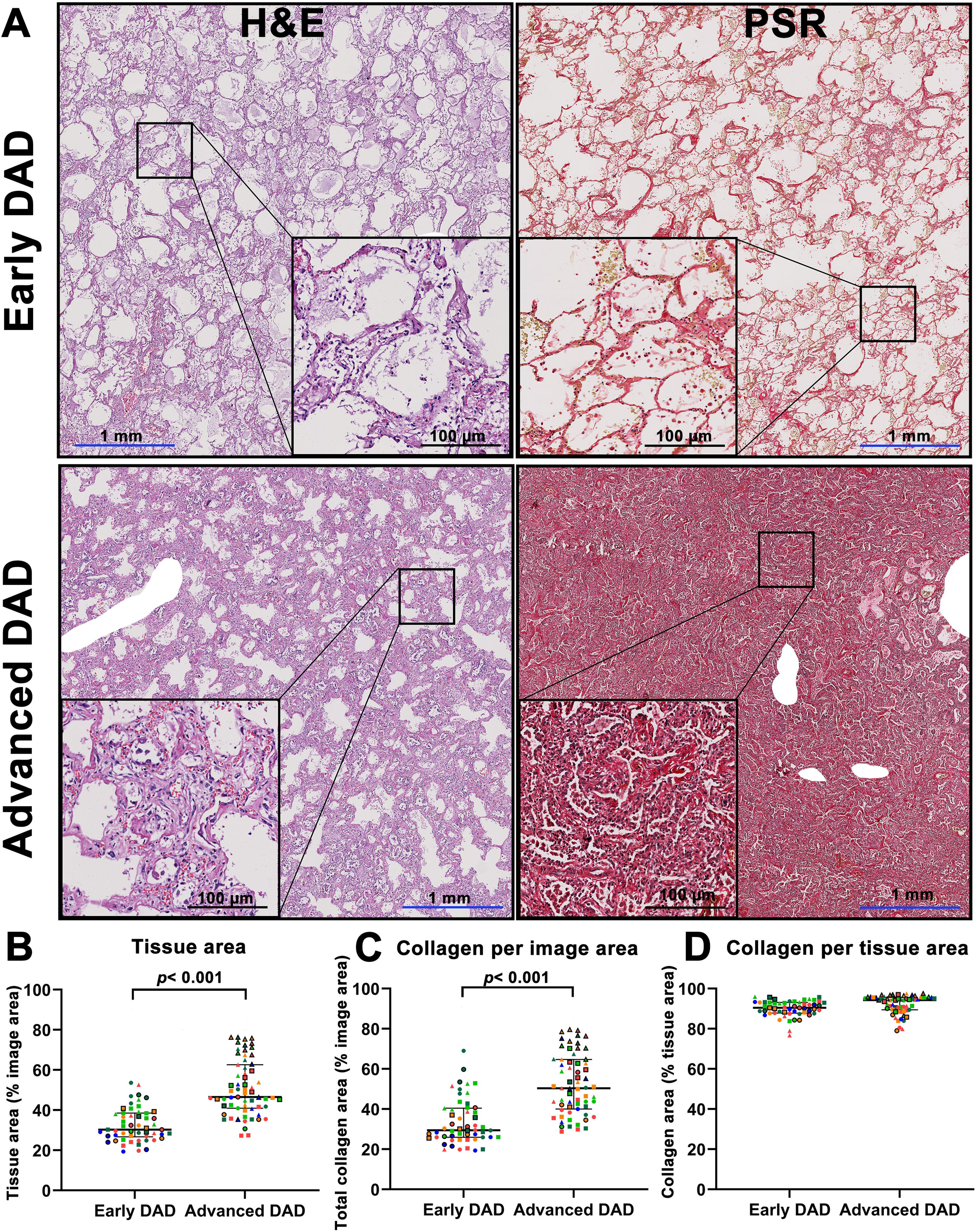
Higher proportional tissue area in the lung regions with advanced DAD. Representative images of parenchymal areas stained with H&E (left column) and PSR (right column) in early and advanced DAD stages (A), showing lung tissue and total collagen, respectively. Images were digitally analyzed to calculate the percentage area within the tissue section perimeter (image area) covered by tissue (B), the percentage of the image area occupied by collagen (C) and the percentage of the tissue area occupied by total collagen (D). Symbols represent individual regions, with those of the same shape and color being from the same patient. Circles represent patients with an ARDS duration <7 days, triangles represent patients with an ARDS duration 7-14 days, and squares represent patients with an ARDS duration >14 days. A linear mixed model was used to test for differences, with early DAD as the reference group adjusted for age and sex. Data are presented as medians and interquartile ranges. P<0.05 was considered significant DAD, Diffuse Alveolar Damage; H&E, hematoxylin and eosin; PSR, Picrosirius Red; ARDS, Acute Respiratory Distress Syndrome.

The percentage of the area within the tissue section perimeter (encompassing both the lung tissue and airspaces inside the outer boundary of the section, but excluding any areas removed for artifacts) that was positive in H&E staining, representing tissue occupancy in the lung samples, was higher in the advanced DAD group (46.5 [41.0-62.6] %) than in the early DAD group (30.3 [26.7-38.5] %), with a *p*-value of 0.0024.

In the PSR staining, where all forms of collagen were detected, while there were differences in the area of the image occupied by collagen, there were no proportional differences in the percentage of tissue occupied by collagen detected. This was evident when assessing both the percentage area and mean intensity of total collagen in the tissue (Figure 3D and Supplemental figure S2A).

### ECM proteins were generally present in the lung parenchyma in ARDS

Having observed a higher proportion of the image occupied by tissue in the advanced DAD stage compared to the early DAD stage, we were interested in further characterizing the origin of this higher proportion of tissue. Further examination of H&E and MTS staining revealed exudative changes, such as intra-alveolar edema and inflammatory cell infiltration, which were predominantly observed in the regions in the early DAD stage. Furthermore, fibrotic alterations, characterized by ECM deposition within the alveolar septa, were predominantly evident in regions with an advanced DAD stage.

Given the observation of fibroproliferative changes in our study and those reported by other studies, we were interested in exploring further details of which ECM elements were participating. Building on previous research results and our focus (33, 34), six ECM proteins were carefully selected for further staining and analysis in lung tissue (Col IIIα1, Col IVα3, Col VIα1, PSTN, LUM, and FN), with their selection guided by their involvement in fibroproliferative processes in lung diseases (18, 35, 36).

All proteins selected, except LUM, were present throughout the parenchyma (figure 4). In 9 of 118 parenchymal regions, over 10% of the area had a LUM-positive signal. Col VIα1 staining showed a particular pattern in the advanced DAD regions, in which the fibers aggregated to form high-intensity bundles (figure 4). Representative images of the staining patterns of ECM proteins in early and advanced DAD are shown in figure 4.

**Figure 4.**
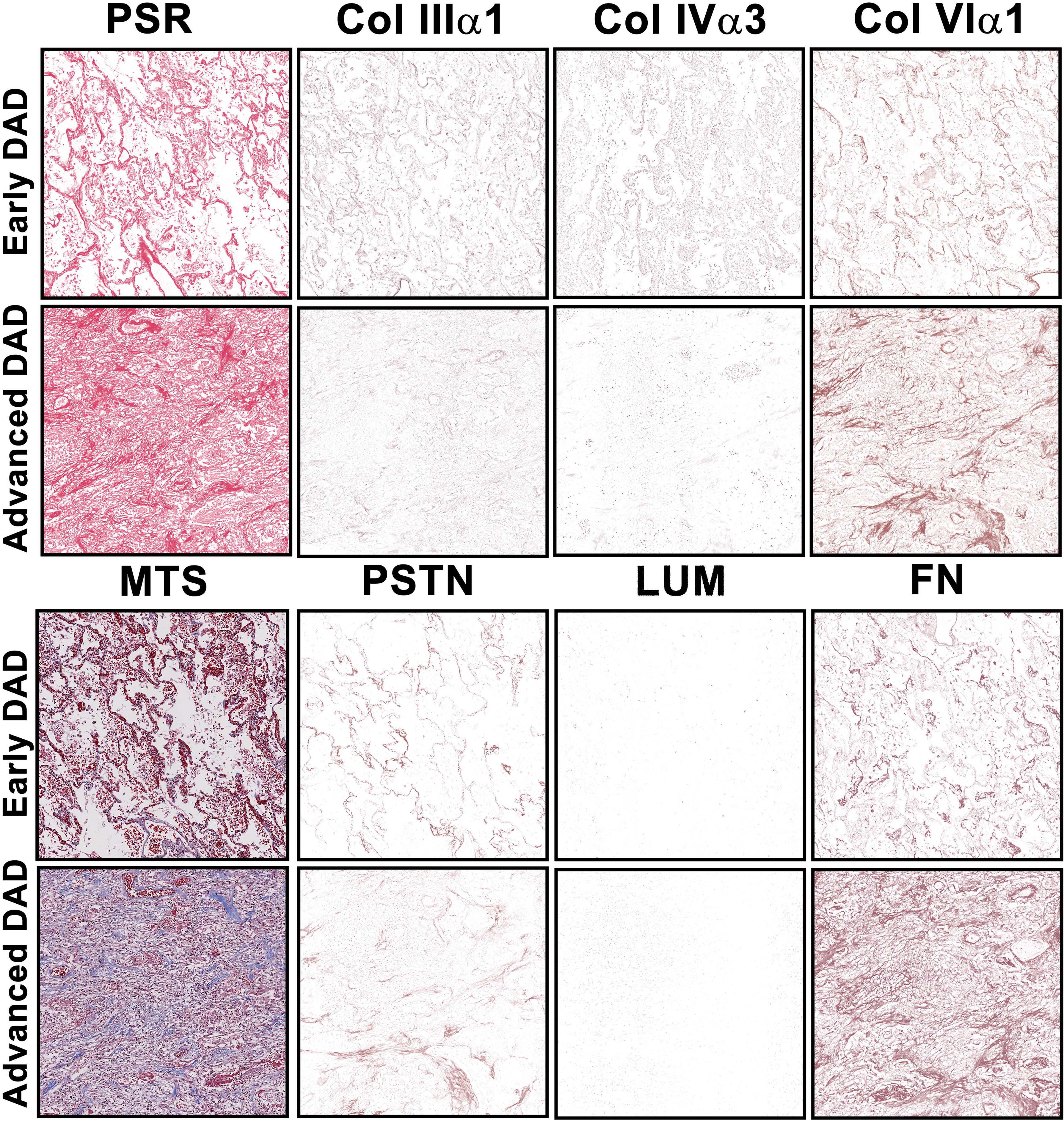
Representative images of ECM protein expression in ARDS lung sections in different pathological stages. Representative images of lung tissues sections with early and advanced stages of DAD stained for PSR to show total collagen deposition and MTS to show collagen and elastin deposition. Addition images illustrate immunohistochemical staining ECM proteins visualized using NovaRed (brown signal). All images were captured at the same location in serial sections from the two regions exemplifying the early and advanced DAD stages. Except for MTS, all images shown are color deconvoluted to display the NovaRed signal alone. DAD, diffuse alveolar damage; MTS, Masson’s trichrome staining; PSR, Picrosirius Red staining; Col IIIα1, collagen type III α1 chain; Col IVα3, collagen type IV α3 chain; Col VIα1, collagen type VI α1 chain; PSTN, periostin; LUM, lumican; FN, fibronectin.

To examine if the effect of the intra-alveolar fibroblasts and intra-alveolar ECM, which were observed in OP, influenced our measurements, we divided the early/advanced DAD each into two subgroups: with or without OP. Since we found there were no differences in the quantification between the subgroups (with/without OP), they were subsequently combined for further calculation and data presentation (data not shown). To show the pathological differences in detail, we have provided a full overview of representative images (original and deconvoluted) of the ECM proteins in early DAD (with or without OP) and advanced DAD (with or without OP), shown in Supplemental figures S3 and S4. A summary of the location of the ECM proteins we investigated in ARDS lung tissues is provided in Supplemental Table S2.

### Lower proportional area of Col IIIα1, Col IVα3, and LUM in the lung regions in advanced DAD stage

To further explore the characteristics of the altered ECM in advanced DAD in ARDS, we examined the proportional differences in the ECM proteins of interest. The proportional area we report represents the tissue area percentage covered by the ECM protein being examined (figure 5). The proportional area of Col IIIα1 was lower in the advanced DAD stage than in the early DAD stage (31.1 [20.5-41.3] vs. 41.4 [29.5-50.2], %, *p*=0.005). A similar difference was observed in Col IVα3 staining, with a lower proportional area in the advanced DAD regions than in the early DAD regions (35.2 [23.6, 49.2] vs. 47.4 [35.6, 56.4], %, *p*=0.002). For LUM, there was also a significantly lower proportional area present in the advanced DAD regions (3.5 [2.3-5.7] vs 4.7 [3.1-7.0], %. *p*=0.002). The proportional tissue area occupied by the remaining ECM proteins (Col VIα1, PSTN, and FN) examined in this study did not differ between the early and advanced DAD regions (figure 5C, D, and F).

**Figure 5.**
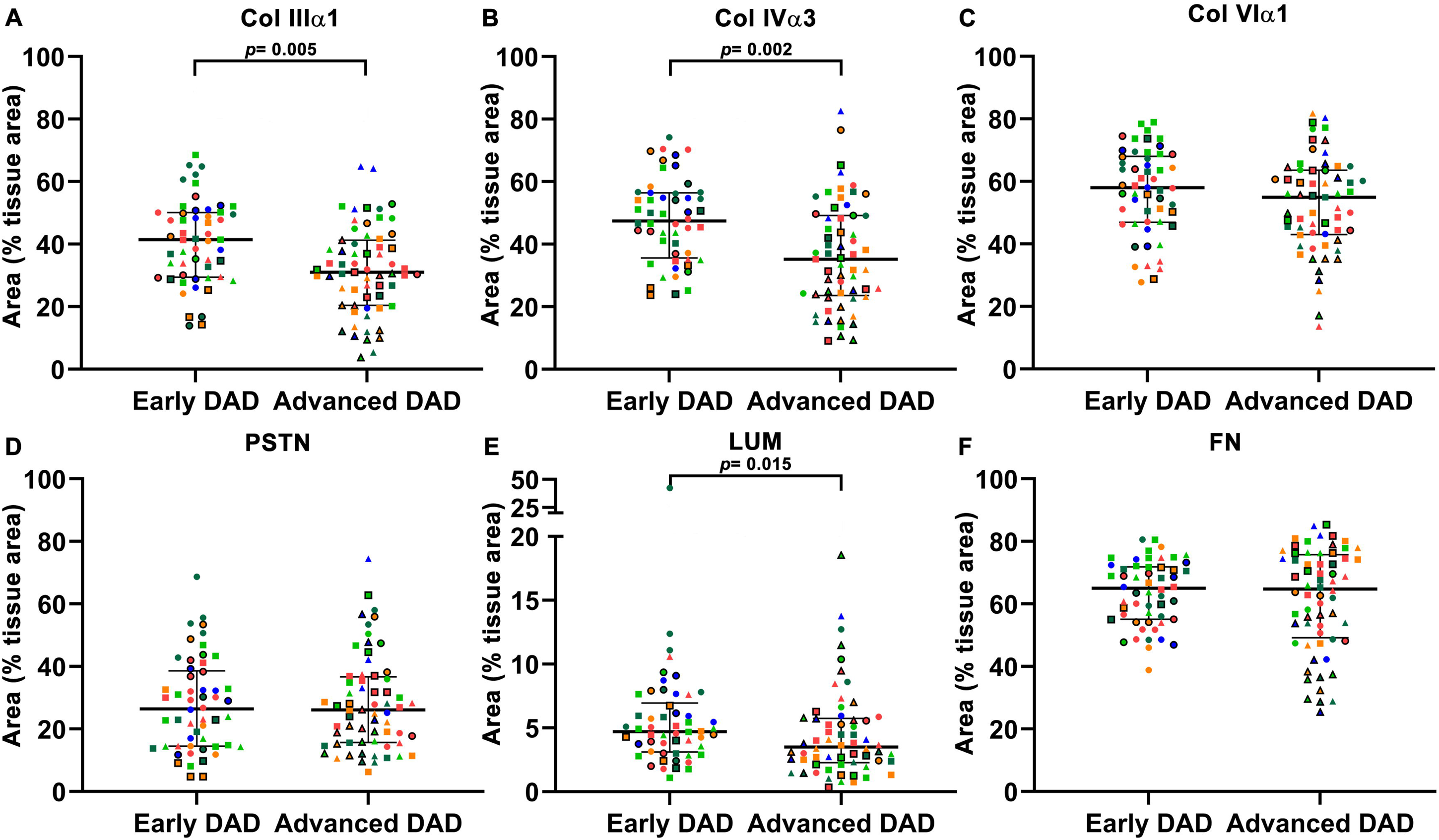
Lower proportional area of Col IIIα regions 1, Col IVα3, and LUM in advanced DAD. The percentage of tissue covered by Col IIIα1 (A), Col IVα3 (B), Col VIα1 (C), PSTN (D), LUM (E) and FN (F), respectively. Symbols represent individual regions, with those of the same shape and color being from the same patient. Circles represent patients with an ARDS duration <7 days, triangles represent patients with an ARDS duration 7-14 days, and squares represent patients with an ARDS duration >14 days. A linear mixed model was used, with early DAD as the reference group adjusted for age and sex. p < 0.05 was considered significant. Ln transformed area data were used for LUM. Data are presented as median and interquartile range. ECM, Extracellular Matrix; Col IIIα1, collagen type III α1 chain; Col IVα3, collagen type IV α3 chain; Col VIα1, collagen type VI α1 chain; PSTN, periostin; LUM, lumican; FN, fibronectin. DAD, Diffuse Alveolar Damage; ARDS, Acute Respiratory Distress Syndrome.

When we examined the average intensity of the signal detected for our ECM proteins, we observed that the Col IIIα1 and Col IVα3 intensities were also lower in the regions with advanced DAD stage than in the early DAD stage (Supplemental figure S2). The intensity signals for the other four ECM proteins (LUM, Col VIα1, PSTN, and FN) did not differ between the DAD stages.

#### A higher density of Col VIα1 bundles per unit of tissue was observed in advanced DAD regions

In the lung parenchyma, Col VIα1 fibers were observed to aggregate into high intensity, bundle-like patterns in the regions of advanced DAD (Figure 6A). The number of Col VIα1 bundles within a given area of tissue was greater in regions from the advanced DAD group, compared to the early DAD group (2.9 [1.6-5.4] vs 1.9 [0.9-3.4], per mm^2, *p*=0.039). However, the average area per bundle (8305.8 [6773.6-11355.0] vs 7151.4 [6087.7-7661.1], μm^2) was similar between the regions of the advanced DAD and the early DAD groups.

**Figure 6.**
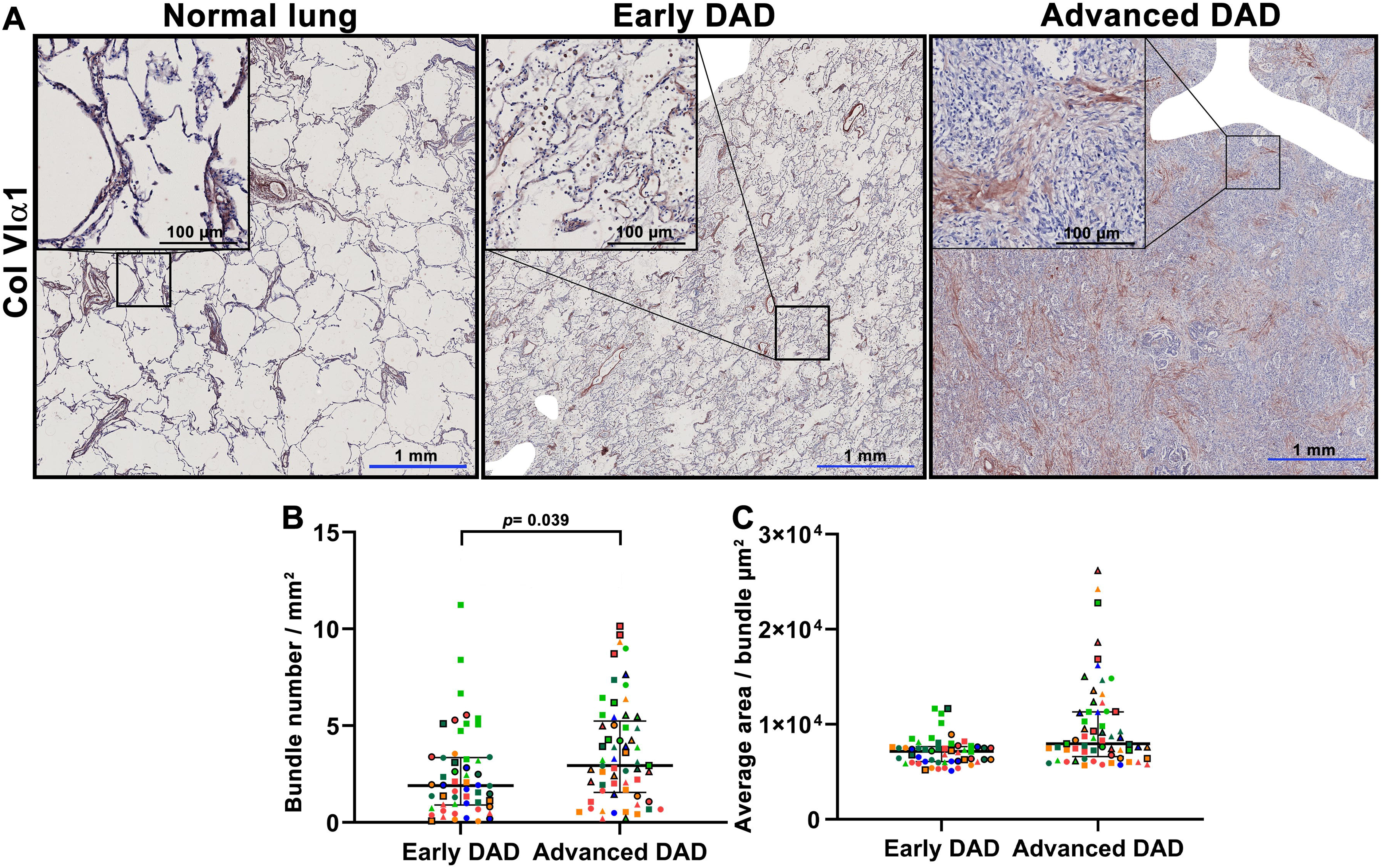
Higher number of Col Viα1 bundles per unit area present in regions with advanced DAD. Representative images of Col VIα1 fiber distribution (detected using Nova Red staining - brown) in normal lung tissue, a region at early DAD stage and a region with advanced DAD (A). The number of Col VIα1 bundles per square millimeter (B) and the average area per bundle were quantified using CellProfiler. Symbols represent individual regions, with those of the same shape and color being from the same patient. Circles represent patients with an ARDS duration <7 days, triangles represent patients with an ARDS duration 7-14 days, and squares represent patients with an ARDS duration >14 days. A linear mixed model was used, with early DAD as the reference group adjusted for age and sex. p < 0.05 was considered significant. Data are presented as median and interquartile range Col VIα1, collagen type VI α1 chain.

### Higher proportional area of total collagen and FN, with lower proportional area of Col IIIα1, Col IVα3 and LUM in lung regions in the extended duration ARDS patient group

Next, we grouped the lung tissue regions into ARDS duration groups to better understand the relationship between the ECM changes in the tissues and the clinical duration of ARDS (figure 7 and Supplemental figure S5).

**Figure 7.**
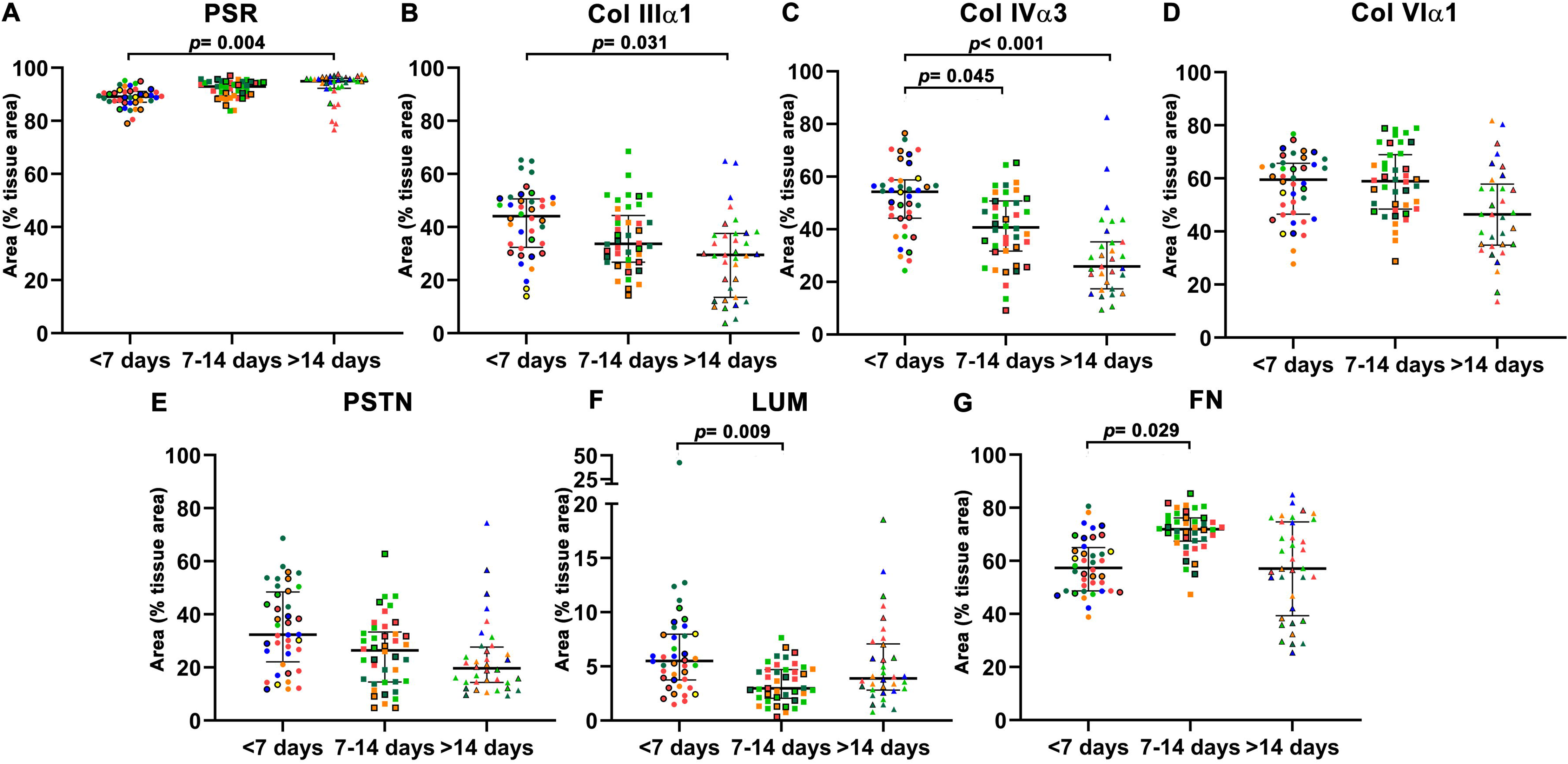
Higher proportional area of total collagen and FN, with lower proportional area of Col IIIα1, Col Ivα 3 and LUM in lung regions from patients with ARDS of longer duration. The percentage of tissue covered by total collagen (PSR) (A), Col IIIα1 (B), Col IVα3 (C), Col VIα1 (D), PSTN (E), LUM (F) and FN (G), respectively. Symbols represent individual regions, with the same shape and color being from the same patient. A linear mixed model was used, with ARDS duration <7 days as the reference group adjusting for age and sex. p < 0.05 was considered significant. Ln transformed area data was used in the LUM dataset. Data are presented as median and interquartile range. ECM, Extracellular Matrix; Col IIIα1, collagen type III α1 chain; Col IVα3, collagen type IV α3 chain; Col VIα1, collagen type VI α1 chain; PSTN, periostin; LUM, lumican; FN, fibronectin; PSR, Picrosirius Red; ARDS, Acute Respiratory Distress Syndrome.

The proportion of total collagen area was higher in the regions from patients with ARDS >14 days, compared to the regions from patients with <7 days of ARDS (95.0 [92.3-96.1] vs 89.1 [87.1-91.1], *p*=0.004). Comparably, the proportional area of FN was significantly higher in the patient group at 7-14 days, compared to the <7 days group (72.0 [85.1-97.5] vs. 57.4 [48.7-65.0], *p*=0.029). However, the proportion of Col IIIα1 (29.6 [13.5-37.7] vs. 44.1 [32.4-50.6], *p*=0.031) and Col IVα3 (25.9 [17.4-35.2] vs. 54.3 [44.2-58.8], *p*<0.001) was significantly lower in the lung regions in the >14 days group than in the <7 days group, and Col IVα3 (40.7 [31.7-50.8] vs 54.3 [44.2-58.8], *p*=0.045) and LUM (3.0 [2.0-4.7] vs 5.5 [3.8-8.0], *p*=0.009) showed significantly lower proportional areas in regions from the 7-14 days group, compared to the <7 days group.

Differences were also observed in the mean intensity of staining across the ARDS duration groups (Supplemental figure S5). The mean intensity of PSR was higher in the 7-14 days and >14 days groups than in the <7 days group, while the mean intensity of Col IIIα1 and PSTN was lower in groups with ARDS duration longer than 7 days than in the <7 days group. The mean intensity of the other ECM proteins examined did not differ among the three ARDS groups.

Interestingly, when grouped by ARDS duration, there was no statistical difference in the number of Col VIα1 fiber bundles per unit area between the groups (Supplemental figure S6). However, the average area of bundles in regions from patients with ARDS duration of >14 days was greater than in the group with <7 days of ARDS duration (8739.0 [7074.7-13028.4] vs 6340.3 [5941.9-7513.8], μm^2, *p*=0.005) (Supplemental figure S6).

## Discussion

In this study, we explored the compositional differences in selected ECM proteins in the lung parenchyma during different stages of DAD. Pathologically, in the advanced DAD stage, the tissue density was higher, while the proportion of collagen type IIIα1 chain, collagen type IVα3 chain and lumican were lower in the lung tissue parenchyma, compared to regions in the early DAD stage. Notably, the organization of collagen type VIα1 chain fibers was denser and more highly concentrated in the regions of advanced DAD. Meanwhile, when considering the clinical duration of ARDS, we found the proportion of tissue occupied by total collagen was greater, as was the proportion of fibronectin present in the lung regions from patients with ARDS duration >7 days groups, compared to those with ARDS of <7 days. Our results indicate that both the composition and organization of the ECM were remodeled in advanced DAD and when patients had ARDS for a longer duration.

Studies investigating the proportional differences in ECM proteins between early and advanced DAD, as occurs during ARDS progression have not been reported before. Moreover, there have not been many studies investigating specific details with respect to the ECM, such as composition and organizational changes in diseased tissues, and ECM differences between different time points. Therefore, our study provides new insights about the contribution of differences in lung ECM protein proportions in different stages of DAD to the pathophysiology of ARDS.

Fibrotic and fibroproliferative changes leading to increase in connective tissue with excessive ECM deposition in ARDS lungs have been previously reported (11, 23, 37, 38). Negri et al. quantified total collagen and elastic fiber density by measuring the stained fiber area per septum in the human lung interstitium (39). They defined early and late ARDS through the presence of histopathological features observed in lung tissues. In the late ARDS group, the lungs showed both higher total collagen and elastin density, compared to early ARDS individuals. However, in contrast to our study, they did not provide information on the type of collagen and other ECM constituents, and they only measured and observed a relatively small sample of lung tissues. A quantitative study, limited to four patients with ARDS, measured the content of collagen I and collagen III in lung autopsies, comparing the results to non-ARDS patients, both with or without mechanical ventilation. The Cyanogen bromide peptide mapping technique was used to study the ratio of type I to type III collagen (40). The higher ratio of collagen type I to type III in ARDS patients suggests that the ECM proportional composition changes in patients with ARDS, and differs from that in normal lung tissue (41).

Proportionally lower collagen type III, type IV and LUM were found in the advanced pathological stage of DAD and also in the tissues from patients with longer duration ARDS. Collagen type III is a fibrillar collagen, and contributes, alongside collagen type I, as one of the major components for forming the structural framework in lung tissue (42, 43). Compared to collagen I, collagen III is less stiff and more flexible (44), therefore lower proportional collagen type III in ARDS lung tissue could indicate a stiffer tissue environment with less compliance (45). Collagen type IV is a network-forming collagen, which is fundamental for maintaining the strength and flexibility of basement membranes, and in this respect is also important for regulating cellular behaviors such as cell-adhesion, signaling and survival (46–49). A lower proportion of collagen type IV in advanced DAD lung tissue could contribute to impaired barrier function of basement membranes, leading to vascular permeability and alveolar edema in ARDS (50). LUM is a small leucine-rich proteoglycan (SLRP) that belongs to the class II subfamily of SLRPs, and locates mainly in the lung interstitium, basement membrane and vessel walls (51). Besides its role as the regulator of collagen fibrillogenesis contributing to the structural integrity of the ECM (52), LUM has been suggested to play a role in promoting epithelial repair processes through interactions with transforming growth factor-b receptor 1 (53). Injured corneal epithelial cells transiently express LUM, suggesting LUM functions to directly regulate epithelial cell migration or adhesion. Compared to controls, LUM-deficient mice have delayed healing of corneal epithelial injury supporting collagen fibrillogenesis independent functions of LUM (54). In our study, the lower proportion of LUM in the lung tissue might indicate a reduced capacity for epithelial repair, potentially contributing to persistent lung injury and impaired tissue recovery during the advanced stage of DAD.

Regulated spatial arrangement and organization of collagen fibers are important for maintaining the ECM structure and function to maintain the stability and functionality of lung tissue (55). Abnormal crosslinking of collagen fibers leads to stiffer tissue and affects cell behavior (56). Under homeostatic conditions, collagen type VI is known to lie along the alveolar wall, functioning as a linking and anchoring element between the basement membrane and interstitial area (57, 58). Collagen type VI α1 deficient mice have decreased dynamic compliance, but increased whole lung and tissue elastance. Their total lung dynamic resistance is increased, whereas the airway resistance is not changed, compared to heterozygous littermates (59). In our study, we observed that in addition to being proportionally lower, Col VIα1 also showed an abnormally dense and highly concentrated fiber organizational pattern in the lung parenchyma; more bundles were observed in advanced, compared to early, DAD regions, and larger bundles were observed in the lungs of patients with ARDS of longer duration, suggesting that the abnormally distributed collagen type VI might play an active role in ARDS progression. For instance, the alterations in collagen type VI might participate in the thickening of the alveolar wall, which results in impaired gas exchange and loss of lung compliance in patients with ARDS.

As a “symptom-defined” syndrome, ARDS has a high reputation of heterogeneity in both clinical and biological studies (60, 61). In our study, we also observed that even within the same lung section, different pathological stages were present in the lungs of the same patient. Additionally, since ARDS evolves as a continuous process, despite our efforts to isolate regions representing distinct pathological stages, some “mixed” areas remained that we were unable to completely separate. Nevertheless, the observation of “mixed” pathological stages within a small area suggests that ongoing damage is likely to have occurred, leading to continuous and repetitive remodeling in the same region. In ARDS, the ongoing insults causing repeated damage could be the continuous force provided by mechanical ventilation and non-resolving inflammation (62–64), which may be drivers of persistent ARDS.

Chronological changes in the histological features of DAD in ARDS lung tissue have been previously reported, indicating that proliferative and fibrotic changes appear as late events (62). Open lung biopsy results from Li et al. also revealed that a longer ARDS duration was independently associated with a higher incidence of fibrosis in patients with ARDS (64). However, Marshall et al. reported early lung fibroproliferative responses, involving biomarkers of collagen remodeling in both BALF and serum, within 24 h of ARDS diagnosis that were associated with mortality (65). Our results in the current study are consistent with previous research. We demonstrated not only an association between longer ARDS duration and a higher number of lung tissue regions in a given area in the advanced DAD stage, but also found that the regions of advanced DAD stage were present in the lung tissues from patients with ARDS duration of less than 7 days.

Our study uncovered novel compositional and organizational changes in ECM proteins during ARDS progression. However, this study had some limitations that can be considered for follow-up research. As a pilot study utilizing human samples, we specifically selected patients who met both the clinical and pathological criteria for ARDS, observed in only half of ARDS cases (6, 66–68), While this resulted in a relatively small sample size, our selection strategy enriched the cohort with patients requiring extended mechanical ventilation and hospitalization, potentially making them more responsive to timely medical interventions (7, 66, 69). Second, due to the challenges in obtaining biopsies, we utilized autopsy and post-transplant lung tissue from patients with non-resolving ARDS. This approach allowed access to multiple regions of the lung, providing a more comprehensive representation of the actual tissue remodeling changes occurring in the lungs. Given that multiple organ failure, rather than respiratory injury, is the leading cause of death in ARDS patients (70), autopsies obtained in our study offer valuable insights into disease progression relatively independent of ARDS severity. Unlike omics data, our focus was not to provide a comprehensive overview of all ECM protein changes in the lung. Instead, we offer a novel perspective by highlighting proportional changes in specific ECM components during ARDS progression. Lastly, due to the pilot nature of the study and the relatively small sample size, we did not incorporate extensive clinical details such as ventilation parameters and medication. However, in statistical evaluation, we accounted for key factors such as age and sex, which have been shown in previous studies to influence proportional changes in ECM (33).

### Conclusion

Our research provides detailed information about proportional differences of specific ECM components in early and advanced DAD, also comparing different duration of ARDS, using human lung tissue. Additionally, we report altered spatial organization of Col VIα1 fibers in the advanced stage of DAD. Our findings underscore the dynamics of ECM remodeling in ARDS development and provide insights into the pathophysiology of ARDS. More translational, mechanistic future studies might help to understand the functional impact and reversibility of the observed ECM changes during ARDS.

## DATA AVAILABILITY

Data is available upon reasonable request from the corresponding author.

## ETHICS DECLARATIONS AND CONSENT TO PARTICIPATE

The study was conducted in accordance with the Research Code of the University Medical Centre Groningen (UMCG), as stated on https://umcgresearch.org/w/research-code-umcg as well as national ethical and professional guidelines Code of Conduct for Health Research (https://www.coreon.org/wp-content/uploads/2023/06/Code-of-Conduct-for-Health-Research-2022.pdf). The use of left-over lung tissue in this study was not subject to Medical Research Human Subjects Act in the Netherlands, as confirmed by a statement of the Medical Ethical Committee of the University Medical Centre Groningen and exempt from consent according to national laws (Dutch laws: Medical Treatment Agreement Act (WGBO) art 458 / GDPR art 9/ UAVG art 24). All donor material and clinical information were deidentified prior to experimental procedures, blinding any identifiable information to the investigators.

## SUPPLEMENTAL MATERIAL

Supplemental Material: Supplemental figure S1-6 and Supplemental table S1-2: DOI ??

## AUTHOR CONTRIBUTIONS

J.K.B., J.P., M.M., J.M. and Y.W.F. conceived and designed research; R.M.J., T. B. performed experiments; Y.W.F, W.T., J.M.V., M.M. analyzed data; J.K.B., J.P., M.M., J.M., W.T., and Y.W.F. interpreted results of experiments; Y.W.F., J.K.B., J.P., M.M., J.M. prepared figures; Y.W.F. J.K.B., J.P., M.M., and J.M. drafted manuscript; Y.W.F. J.K.B., J.P., M.M., J.M. R.M.J., T.B., J.M.V., and W.T. edited and revised manuscript; Y.W.F. J.K.B., J.P., M.M., J.M. R.M.J., T.B., J.M.V., and W.T. approved final version of manuscript.

## Supporting information

publication licence for figure 2

