## Supplemental material for "Compositional changes of the lung extracellular matrix in acute respiratory distress syndrome"

RESEARCH ARTICLE

running head: compositional changes of ecm protein in ards

*Contributed equally

Correspondence:

Janesh Pillay

**Supplemental data Appendix**

**Contents**

**Supplemental table S1 Detailed information of antibodies used for immunohistochemical staining of ECM proteins	3**

**Supplemental Material S1: Macro template used in the data generation in Fiji	4**

**Supplemental figure S1 Schematic representation of the process for the detection of Col VIα1 bundles using CellProfiler.	6**

**Supplemental figure S2: Mean intensity of detected ECM proteins in regions of early and advanced DAD	7**

**Supplemental figure S3 Representative staining images of ECM proteins in different DAD pathological stages in lung sections of ARDS patients (original staining image)	8**

**Supplemental figure S4 Representative staining images of ECM proteins in different DAD pathological stages in lung sections of ARDS patients (deconvoluted image)	9**

**Supplemental table S2. The overview of ECM proteins expression in different locations in ARDS lung tissue	10**

**Supplemental figure S5: Higher mean intensity of total collagen, but lower mean intensity of Col IIIα1, PSTN and LUM in lung regions from patients with longer duration ARDS	11**

**Supplemental figure S6 Col VI bundles with higher average area present in the lung regions from patients with longer duration ARDS**.	**12**

**Supplemental table S1 Detailed information of antibodies used for immunohistochemical staining of ECM proteins**

| **Catalog of Ab (Company)** | **Target** | **Antigen retrieval** | **Primary Ab concentration (host species, clonality)** | **Incubation time and temperature** | **Secondary Ab** | **Tertiary Ab** |
| --- | --- | --- | --- | --- | --- | --- |
| ab7778 (Abcam) | Col IIIα1 | Citrate (10mM, pH 6.0) | 11.6 μg/mL (rabbit, polyclonal) | Overnight incubation, 4°C | HRP conjugated Goat anti-Rabbit (1:100, P0448, Dako, Denmark) | HRP conjugated Rabbit anti-Goat (1:100, P0449, Dako, Denmark) |
| NBP1-68939 (Novus Biologicals) | Col IVα3 | Pepsin/HCl solution (0.03mM pepsin in 0.01M HCl) | 5 μg/mL (rabbit, polyclonal) |  |  |  |
| NB120-6588 (Novus Biologicals) | Col VIα1 | Citrate (10mM, pH 6) | 10 μg/mL (rabbit, polyclonal) |  |  | / |
| ab168348 (Abcam) | LUM |  | 0.2 μg/mL (rabbit, monoclonal) |  |  | / |
| ab14041 (Abcam) | PSTN | Tris/EDTA buffer (10mM, pH 9) | 0.25 μg/mL (rabbit, polyclonal) |  |  | / |
| ab66584 (Abcam) | FN |  | 1 μg/mL (rabbit, polyclonal) | 60min incubation, room temperature |  | HRP conjugated Rabbit anti-Goat (1:100, P0449, Dako, Denmark) |

When performing batch staining procedures that require more than one vial of antibody, we ensured that all vials were sourced from the same production batch. The concentration of Abs was batch dependent.

ECM, extracellular matrix; Ab, antibody; Col IIIα1, collagen III α1; Col IVα3, collagen IVα3; Col VI α1, collagen VI α1;HCl, Hydrochloric Acid; Tris, Tris(hydroxymethyl)aminomethane; EDTA, Ethylenediaminetetraacetic Acid; HRP, Horseradish Peroxidase.

**Supplemental Material S1: Macro template used in the data generation in Fiji**

//This macro performs color deconvolution and measures positive pixels in the deconvoluted images.

//Fill in the name of the deconvolution vector in the highlighted part (optimized vector).

// Optimized vector will separate in colors: blue (hematoxylin), Red (NovaRed), and background (RBCs)

// Set foreground, background, and selection colors

run("Colors...", "foreground=black background=white selection=yellow");

// Get name of image being processed

name=getTitle();

print(name);

// Set measurements needed for analysis

run("Set Measurements...", "area mean modal integrated median skewness kurtosis area_fraction limit display redirect=None decimal=6");

// Run colour deconvolution

run("Colour Deconvolution", "vectors=[*optimized vector*] ");

// Merge blue and red images and measure positive pixels for the total area

imageCalculator("AND create", name+"-(Colour_1)",name+"-(Colour_2)");

selectImage("Result of "+name+"-(Colour_1)");

setAutoThreshold("Default");

setThreshold(*optimized threshold for total area*);

run("Measure");

close();

// Close Image of Colour_1, not needed for analysis

selectWindow(name+"-(Colour_1)");

close();

// Fill in below the thresholds for Colour2 image (total thresholds)

selectWindow(name+"-(Colour_2)");

setAutoThreshold("Default");

setThreshold(*optimized threshold for positive signal*);

run("Measure");

//

The total threshold is divided into 6 separate regions, for calculating the weak, medium, and strong signal area; the numbers below are illustrative when using a total threshold from 0 - 200

setThreshold(168, 200);

run("Measure");

setThreshold(134, 167);

run("Measure");

setThreshold(101, 133);

run("Measure");

setThreshold(67, 100);

run("Measure");

setThreshold(34, 66);

run("Measure");

setThreshold(0, 33);

run("Measure");

run("Close All");

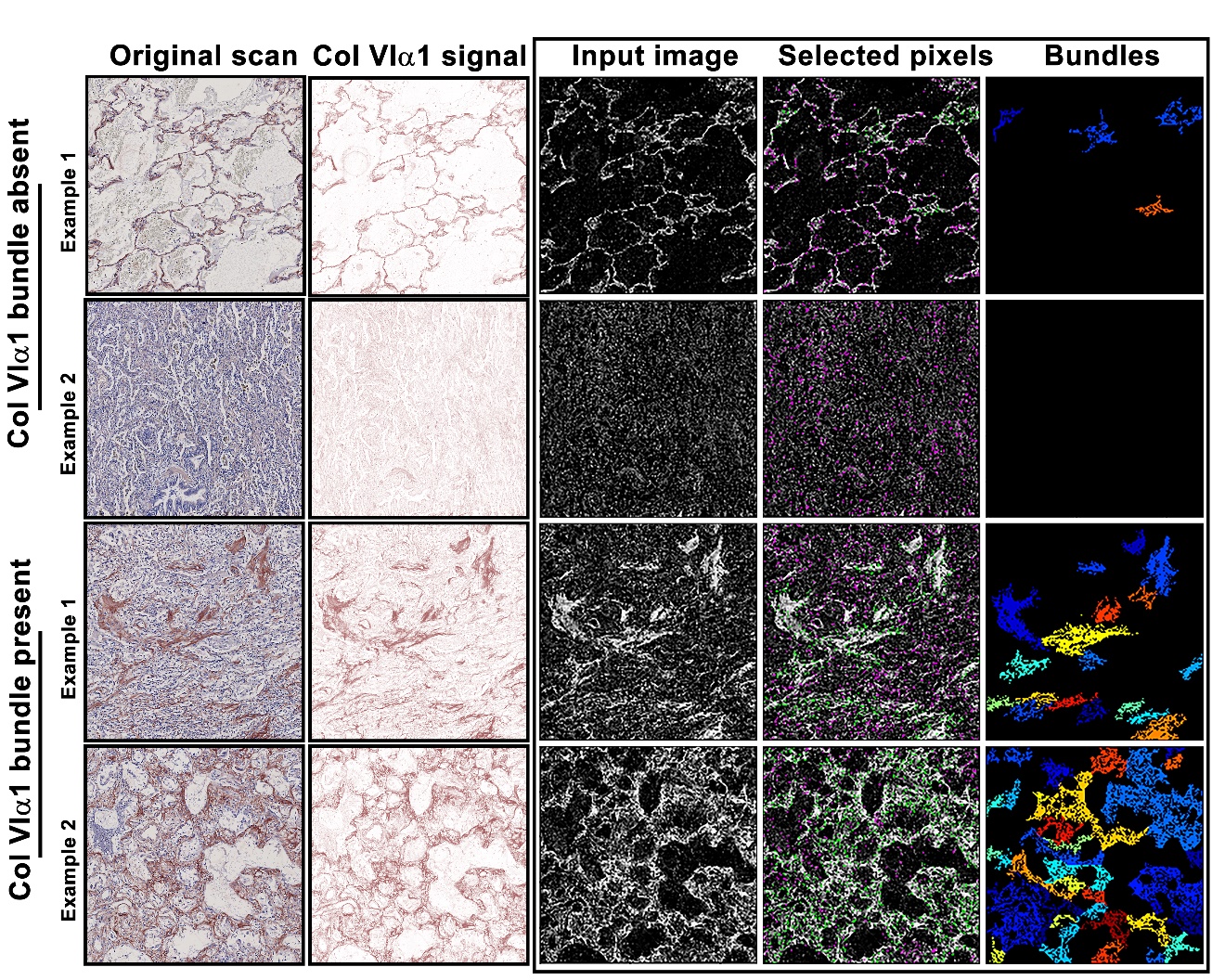

**Supplemental figure S1 Schematic representation of the process for the detection of Col VIα1 bundles using CellProfiler**

Illustration of the process used for detecting Col VIα1 bundles in lung parenchymal images using the image analysis software CellProfiler. The first two rows show examples of images without Col VIα1 bundles present, while the lower two rows show examples of images with condensed Col VIα1 bundles.

The first column shows the original scan images, The second column shows the color deconvoluted images, which represented Col VIα1 signal detected with NovaRed. The third column presents the input images processed with “invert” operation for enhanced contrast to aid in feature detection. The fourth column shows the processed image overlaid with the colored outlines of the identified objects. Each object is assigned one of two (default) colors (Green: Acceptable; Magenta: Discarded based on size). The fifth column displays the identified bundles, color-coded to distinguish separate structures within the tissue.

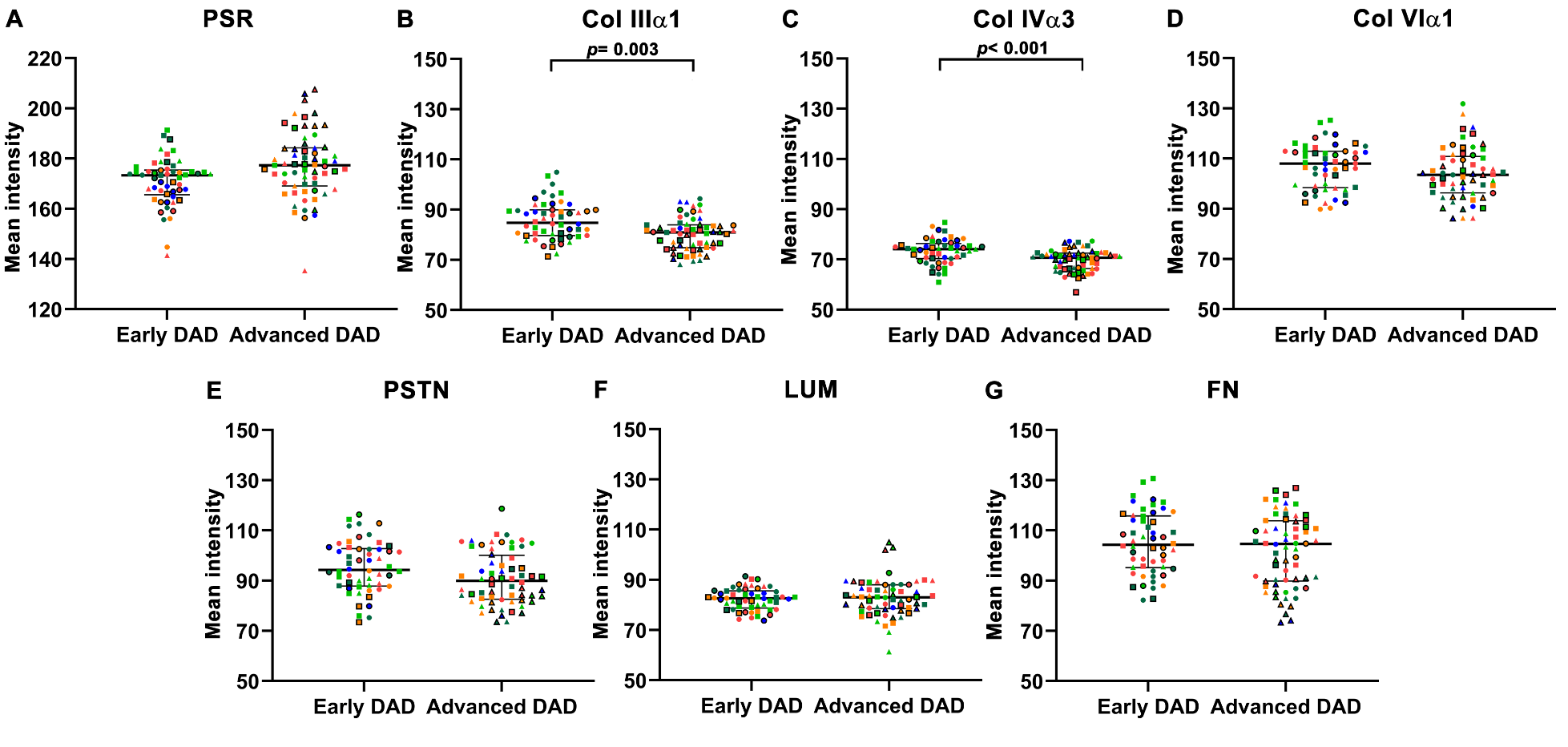

**Supplemental figure S2: Mean intensity of detected ECM proteins in regions of early and advanced DAD**

Images from lung tissue sections, partitioned for early and late DAD stages were digitally analyzed and the mean intensity of pixels positive for the detected protein within the tissue region area was calculated. (A) total collagen (stained with PSR), (B) Col IIIα1, (C) Col IVα3, (D) Col VIα1, (E) PSTN, (F) LUM and (G) FN .

Each data point represents an individual region, with combinations of shapes and colors indicating regions from the same patient. Circle, square and triangle dots represent the regions from patients of ARDS duration <7, 7-14, and >14 days groups, respectively.

A linear mixed model was used for statistics, with the early DAD as reference group, adjusting for age and sex. p<0.05 was considered significant. Ln (normal log) transformed area data was used analyzes of the PSR dataset. Data are presented as median and interquartile range.

ECM, Extracellular matrix; Col IIIα1, collagen type III α1 chain; Col IVα3, collagen type IV α3 chain; Col VIα1, collagen type VI α1 chain; PSTN, periostin; LUM, lumican; FN, fibronectin. DAD, diffuse alveolar damage; ARDS, Acute Respiratory Distress Syndrome.

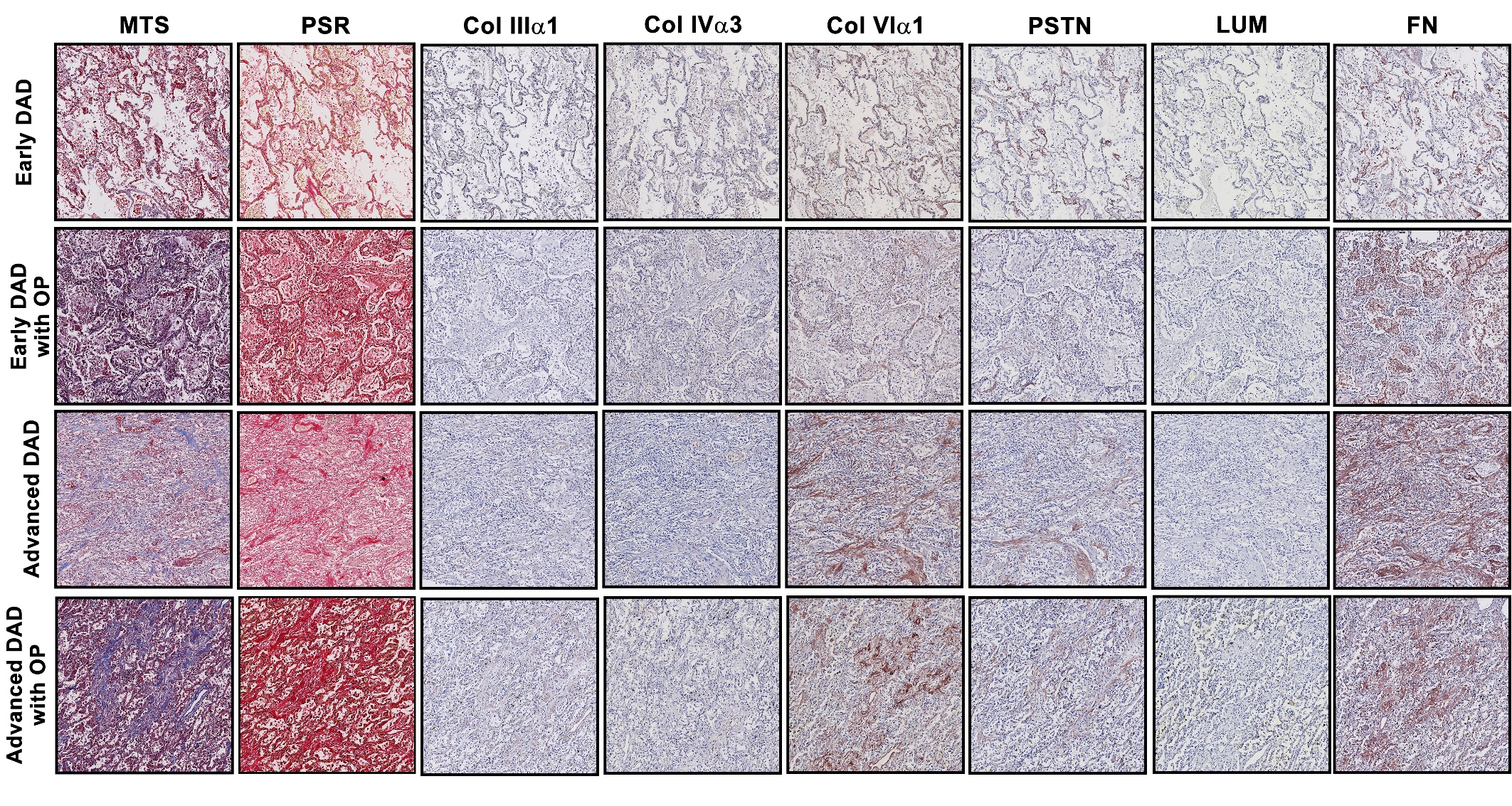

**Supplemental figure S3 Representative staining images of ECM proteins in different DAD pathological** **stages in lung sections of ARDS patients (original staining image)**

Representative photomicrographs of the original scanned images captured at the same location in serial sections of the regions featuringearly DAD, early DAD with OP, advanced DAD and advanced DAD with OP. The stainings presented include histochemical staining (MTS and PSR staining), and immunohistochemical staining for different ECM proteins.

OP, organizing pneumonia; MTS, Masson’s trichrome staining; PSR, Picrosirius Red staining; Col IIIα1, collagen type III α1 chain; Col IVα3, collagen type IV α3 chain; Col VIα1, collagen type VI α1 chain; PSTN, periostin; LUM, lumican; FN, fibronectin.

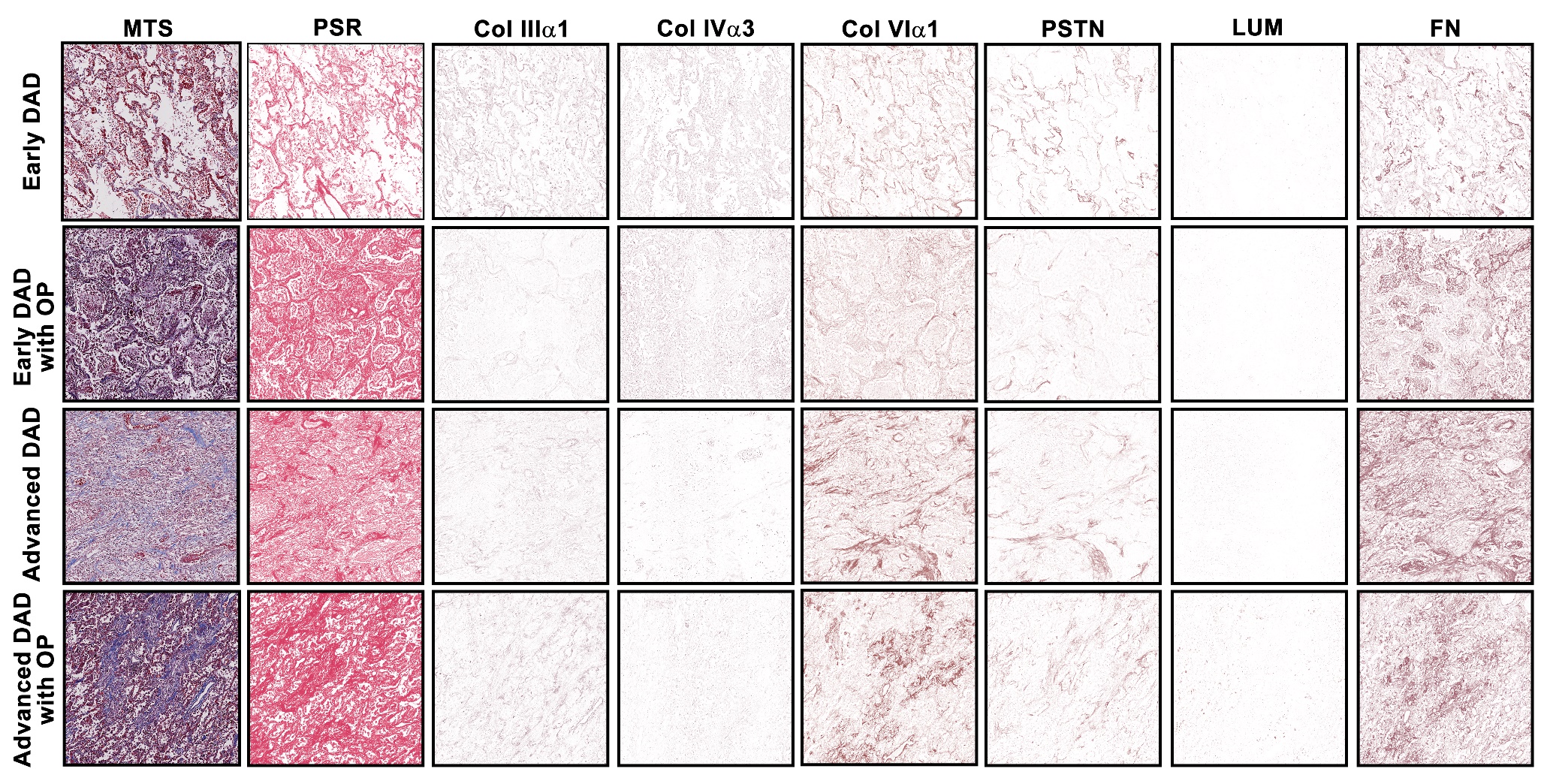

**Supplemental figure S4 Representative staining images of ECM proteins in different DAD pathological stages in lung sections of ARDS patients (deconvoluted image)**

Representative photomicrographs of the original scanned images of MTS, and color deconvoluted images for PSR staining and specific ECM proteins. Except MTS and PSR, the signals of ECM proteins were detected using NovaRed. The representative images show the expression of ECM at the same location in serial sections of the regions featuring early DAD, early DAD with OP, advanced DAD and advanced DAD with OP.

OP, organizing pneumonia; MTS, Masson’s trichrome staining; PSR, Picrosirius Red staining; Col IIIα1, collagen type III α1 chain; Col IVα3, collagen type IV α3 chain; Col VIα1, collagen type VI α1 chain; PSTN, periostin; LUM, lumican; FN, fibronectin.

**Supplemental table S2. The overview of ECM proteins expression in different locations in ARDS lung tissue**

| **ECM protein** | **Parenchyma** | **Airway** | | | **Blood vessel** | |
| --- | --- | --- | --- | --- | --- | --- |
|  |  | **Epithelium** | **BM** | **ASM** | **endothelium** | **VSM** |
| Col IIIα1 | + | +/- | + | + | + | +/- |
| Col IVα3 | + | + | + | + | + | + |
| Col VIα1 | + | - | + | + | + | + |
| PSTN | + | - | + | +/- | +/- | +/- |
| LUM | +/- | + | - | +/- | +/- | +/- |
| FN | + | - | + | + | + | +/- |

ECM, extracellular matrix; BM, basement membrane; ASM, airway smooth muscle; VSM, blood vessel smooth muscle; Col IIIα1, collagen type III α1 chain; Col IVα3, collagen type IV α3 chain; Col VIα1, collagen type VI α1 chain; PSTN, periostin; LUM, lumican; FN, fibronectin; (-) no staining; (+) positive staining; (+/-) variability in strength of positive signal dependent on tissue donor.

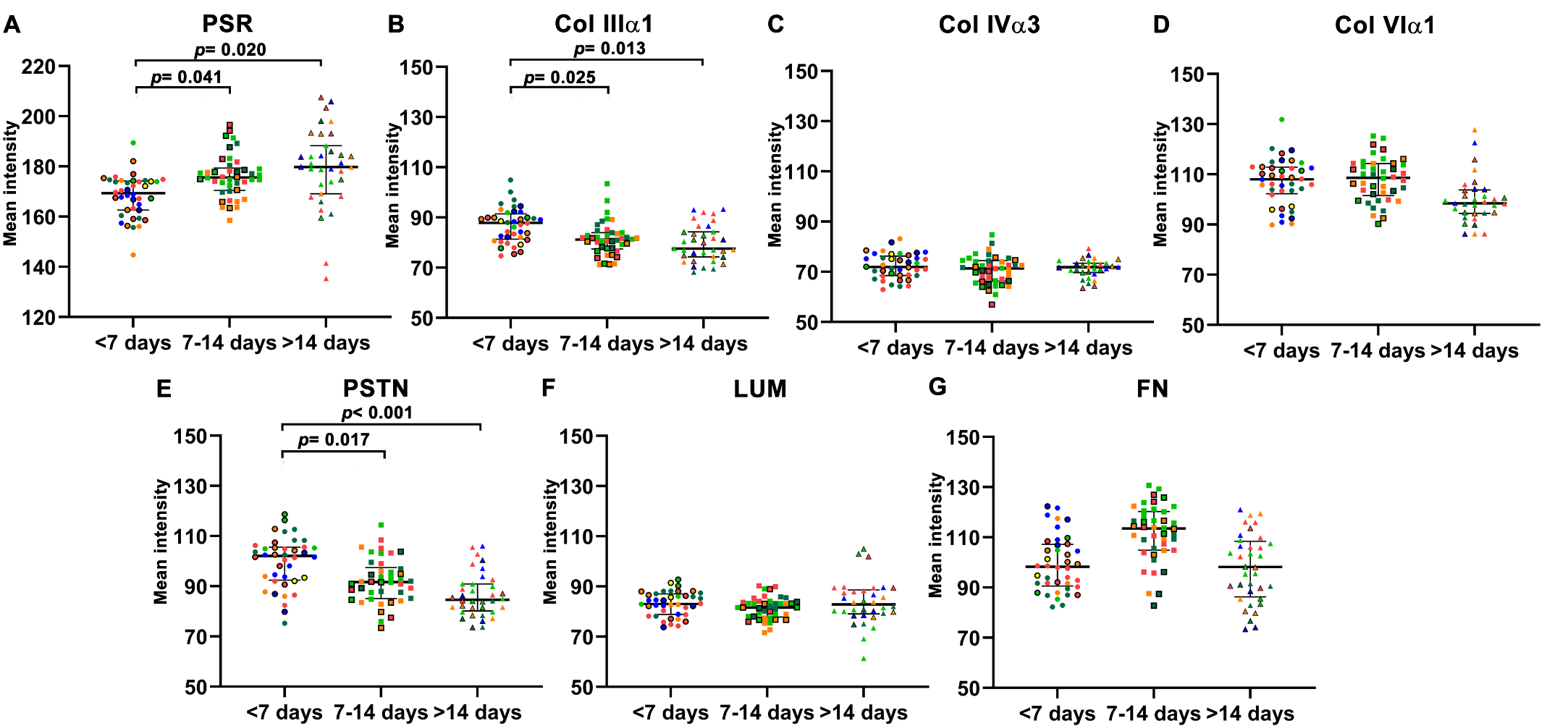

**Supplemental figure S5: Higher mean intensity of total collagen, but lower mean intensity of Col IIIα1, PSTN and LUM in lung regions from patients with longer duration ARDS**

Images from lung tissue sections, partitioned for duration of clinical ARDS were digitally analyzed and the mean intensity of pixels positive for the detected protein within the tissue region area was calculated. (A) total collagen (stained with PSR), (B) Col IIIα1, (C) Col IVα3, (D) Col VIα1, (E) PSTN, (F) LUM and (G) FN. Each data point represents an individual region, with combinations of shapes and colors indicating regions from the same patient. A linear mixed model was used in the statistics, with the <7 days group as the reference group, adjusting for age and sex. p<0.05 was considered as significant. Ln transformed data was used in the PSR dataset in the calculation. Data are presented as median and interquartile range.

ECM, extracellular matrix; Col IIIα1, collagen type III α1 chain; Col IVα3, collagen type IV α3 chain; Col VIα1, collagen type VI α1 chain; PSTN, periostin; LUM, lumican; FN, fibronectin; PSR, Picrosirius Red; ARDS, Acute Respiratory Distress Syndrome.

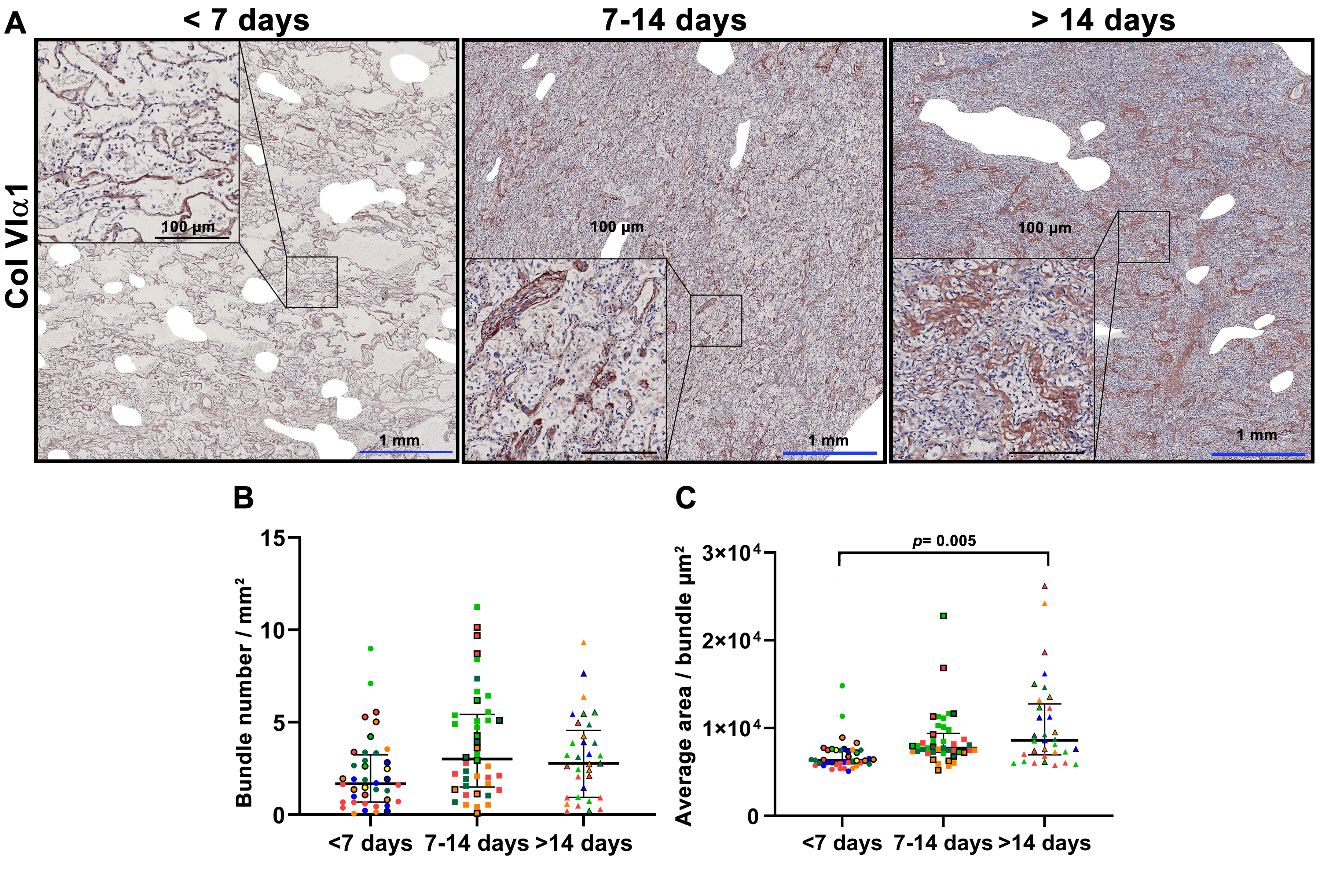

**Supplemental figure S6 Col VI bundles with higher average area present in the lung regions from patients with longer duration ARDS**

Representative images of Col VIα1 fiber distribution (detected using Nova Red staining - brown) lung sections from a patient with ARDS duration <7 days, ARDS duration 7-14 days and ARDS duration >14 days (A). The number of Col VIα1 bundles per square millimeter (B) and the average area per bundle were quantified using CellProfiler. Symbols represent individual regions, with the same shape and color being from the same patient. A linear mixed model was used, with ARDS duration <7 days as the reference group adjusting for age and sex. p < 0.05 was considered significant. Data are presented as median and interquartile range

Col VIα1, collagen type VI α1 chain.
